# Circuit directionality for motivation: lateral accumbens-pallidum, but not pallidum-accumbens, connections regulate motivational attraction to reward cues

**DOI:** 10.1101/474387

**Authors:** Elizabeth B. Smedley, Alyssa DiLeo, Kyle S. Smith

## Abstract

Sign-tracking behavior, in which animals interact with a cue that predicts reward, provides an example of how incentive salience can be attributed to cues and elicit motivation. The nucleus accumbens (NAc) and ventral pallidum (VP) are two regions involved in cue-driven motivation. The VP, and subregions of the NAc including the medial shell and core, are critical for sign-tracking, and connections between the medial shell and VP are known to participate in sign-tracking and other motivated behaviors. The NAc lateral shell (NAcLSh) is a distinct and understudied subdivision of the NAc, and its contribution to the process by which reward cues acquire value remains unclear. The NAcLSh has been implicated in reward-directed behavior, and has reciprocal connections with the VP, suggesting that NAcLSh and VP interactions could be important mechanisms for incentive salience. Here, we use DREADDs (Designer Receptors Exclusively Activated by Designer Drugs) and an intersectional viral delivery strategy to produce a biased inhibition of NAcLSh neurons projecting to the VP, and vice versa. We find that disruption of connections from NAcLSh to VP reduces sign-tracking behavior while not affecting consumption of food rewards. In contrast, VP to NAcLSh disruption affected neither sign-tracking nor reward consumption, but did produce a greater shift in animals’ behavior more towards the reward source when it was available. These findings indicate that the NAcLSh→VP pathway plays an important role in guiding animals towards reward cues, while VP→NAcLSh back-projections may not and may instead bias motivated behavior towards rewards.

## 1 Introduction

Sign-tracking, or autoshaping, includes a behavioral phenomenon where animals interact with a conditioned stimulus (CS+) that predicts an unconditioned stimulus (US), like a reward, even though the US delivery is not contingent on this behavior (Brown and Jenkins 1968; Flagel and Robinson 2017; Boakes 1977). Sign-tracking reflects the attribution of incentive salience to the CS+ and can be highly sensitive to changes in motivational state and cue-reward relationships (Jenkins and Moore 1973; Robinson and Berridge 2013; Berridge and Robinson 2003; Berridge 2004; Chang and Smith 2016; Smedley and Smith 2018a, 2018b; Flagel and Robinson 2017). The nucleus accumbens (NAc) and ventral pallidum (VP), two reciprocally connected limbic regions, have long been implicated in motivated behaviors directed towards CS+s and their paired rewards (Smith et al. 2009; Root et al. 2015; Mogenson 1980). For example, manipulations to decrease the function of the NAc can result in reduced sign-tracking behaviors (Chang and Holland 2012; Chang and Holland 2013; Cardinal et al. 2002). Phasic activity patterns of NAc neurons, and release patterns of neurotransmitter input, can also represent the reward-related value of CS+ cues including those that evoke sign-tracking behavior (Day and Carelli 2007; Day et al. 2006; Batten et al. 2018; Flagel et al. 2011; Singer et al. 2016; Wan and Peoples 2008; Ambroggi et al. 2011). VP neuronal activity is similarly modulated by CS+ cues that are imbued with incentive salience (Tindell 2005; Tindell 2009; Smith et al. 2011; Richard et al. 2016; Ahrens et al. 2016; Ahrens et al. 2018). Inhibition of the VP also disrupts sign-tracking behavior, and does so in a manner not attributable to changes in motor expression or the value of the reward (Chang, Todd, Bucci, & Smith 2015).

The contributions of different anatomically organized projections between the NAc and VP to incentive salience remains unclear. The NAc can be divided into core and shell regions, and the shell further subdivided into medial and lateral segments (Heimer et al. 1997; Zahm 2000; van Dongen et al. 2005; Yang et al. 2018; Zahm and Brog 1992; Kuo and Chang 1992; Zaborsky et al. 1985). There is evidence that connections between the medial NAc shell and VP play an important role in forms of appetitive motivation including cue-triggered reward seeking, reward consumption, and sign-tracking (Stratford and Kelley 1997, Smith and Berridge 2007, Leung and Balleine 2015, Chang et al 2018, Smith et al. 2011).

In contrast, roles for connections between the lateral NAc shell (NAcLSh) and VP remain highly understudied. The NAcLSh itself participates in positive motivation (Zhang and Kelley 2000; Lammel et al. 2012; Mahler and Aston-Jones 2012; Yang et al. 2018). However, it is distinct from medial NAc shell in its neurochemical and anatomical connectivity (Zahm and Brog 1992, Deutch and Cameron 1993). For example, the NAcLSh has reciprocal connections with VP in a more mid-lateral VP zone that is partly dissociable from medial NAc shell-VP connectivity (Brog et al., 1993, Zahm 2000; Churchill and Kalivas, 1994). NAcLSh is also connected with areas linked with appetitive behavior including the ventral tegmental area, substantia nigra pars compacta, lateral hypothalamus, and extended amygdala (Yang et al. 2018, Heimer et al. 1991, Groenewegen and Russchen 1984, Brog et al. 1993). Thus, anatomically, both NAcLSh and VP structures are poised to interact with one another reciprocally and to affect a broader neural network that includes areas implicated in motivation and behavioral control.

To begin addressing the role of this reciprocal connection in motivation, we investigated the effect of biasing chemogenetic inhibition of VP projections to the NAcLSh (VP→NAcLSh), or NAcLSh projections to the VP (NAcLSh→VP), on sign-tracking for food and on the primary motivation to eat food. We found that NAcLSh→VP selectively reduced sign-tracking, that VP→NAcLSh inhibition selectively increased goal-approach that also occurred during the cues, and that neither pathway manipulation detectably affected free feeding behavior. These results highlight a preferential role for the NAcLSh→VP pathway in regulating the motivational attraction to reward-paired cues. Moreover, the functional dissociation between the pathway manipulations indicates that the NAcLSh→VP conveys information to the VP for the regulation of reward cue attraction that could be insensitive to the integrity of information transferred back from the VP.

## 2 Materials and Methods

### 2.1 Subjects

Experimentally naïve male Long Evans rats (arrival weight 250-300g) were obtained from Charles River (n = 52; Charles River, Indianapolis, IN, USA). Rats were single housed in ventilated plastic cages in a climate-controlled colony room set to a 12h light/dark cycle (lights on at 7:00 A.M.). Experiments were conducted during the light cycle. Food and water were available *ad libitum* until 7 days before magazine training, at which point weight was restricted to 85% of *ad libitum* weight prior to testing. Restriction was maintained throughout the experiment. For restriction, rats were provided with 5-12g of standard chow (Harlan Teklad 2014) and free access to water after each testing session. All procedures were approved by the Dartmouth Institutional College Animal Care and Use Committee.

### 2.2 Surgical Procedures

Rats were anesthetized with isoflurane gas and placed in a stereotaxic apparatus (Stoelting, Kiel, WI, USA). Surgery was conducted under aseptic conditions. A 5 µl, 33-gauge beveled needle-tipped syringe (World Precision Instruments, Sarasota, FL, USA) was lowered to the bilateral target sites and allowed to rest for 3 min. Viral vectors were infused at a rate of 0.15 µl/min the following targets in mm from bregma: VP (−0.12 AP, +/−2.4 ML, −8.2 DV), NAcLSh (+1.44 AP, +/−2.0 ML, −8.2 DV) (Paxinos and Watson, 2007).

Post-infusion, the needle was allowed to rest for 5 minutes to all viral dispersion. Two groups of animals were used to assess the NAcLSh→VP pathway: NAcLSh→VP Inhibition and NAcLSh→VP Control. The inhibition group received 0.6 µl of AAV-hSyn-DIO-hM4D(Gi)-mCherry (n = 13; AAV5, UNC Vector Core; n = 4; AAV8, Addgene) in the NAcLSh and 1.0 µl CAV-Cre (IGMM, France) or CAV-Cre-GFP (IGMM, France) in the VP. Controls for the NAcLSh→VP projection received an AAV vector lacking the DREADD molecule (AAV5-hSyn-DIO-mCherry; n = 9, UNC Vector Core) in the NAcLSh and also received an infusion of 1.0 µl CAV-Cre or CAV-Cre-GFP in the VP.

Likewise, two groups of animals were used to assess the VP→NAcLSh pathway: VP→NAcLSh Inhibition and VP→NAcLSh Control. The inhibition group received 0.6 µl of AAV-hSyn-DIO-hM4D(Gi)-mCherry (n = 12, AAV5, UNC Vector Core; n = 4, AAV8, Addgene) in the VP and 1.0 µl CAV-Cre or CAV-Cre-GFP in the NAcLSh. Controls of the VP→NAcLSh projection received the same virus structure with the omission of the receptor (AAV5-hSyn-DIO-mCherry; n = 10, UNC Vector Core) in VP and also received an infusion of 1.0 µl CAV-Cre or CAV-Cre-GFP in the NAcLSh. Surgical incisions were closed with surgical clips and covered with Neosporin. Rats were given IP injections 3 mg/kg of Ketoprofen and 5 ml of 0.9% sterile saline after surgery and monitored for the remainder of the experiment. Clips were removed under isoflurane anesthetic within 2 weeks of surgery. Animals were recovered with food, DietGel (Clear H2O, ME, USA), and water. Food restriction and behavioral procedures began a minimum of 3 wks post-surgery.

### 2.3 Test Apparatus

Sign-tracking procedures were conducted in standard operant chambers (Med Associates, St. Albans, VT) that were enclosed in sound- and light-attenuating cabinets and were outfitted with fans for ventilation and white noise. Chambers contained two retractable levers on either side of a recessed magazine where food rewards would be delivered. Lever depressions were recorded automatically, and magazine entries were recorded through breaks in an infrared beam at the magazine site. Free feeding procedures were conducted in cleaned plastic home-cages affixed with a glass petri dish for containing food.

### 2.4 Test Procedures

The experimental design included one day of magazine training, twelve days of sign-tracking training, and 2 days of free-feeding testing. For magazine training, one 30 min session of magazine training was conducted to habituate rats to the chamber and grain pellet reward delivery (BioServ, 45 mg Dustless Precision Pellets, Rodent Grain-Based Diet). Pellets were delivered such that over a 30 min period about 60 pellets were delivered [*p(pellet per second) = 1/30*]. Magazines were checked after testing to confirm consumption of reward pellets.

Sign-tracking training then began for 12 consecutive daily sessions. A given session contained 25 CS+ trials where the 10 sec insertion of a retractable lever was followed by the delivery of 2 grain pellets into the magazine, and 25 CS-trials where the 10 sec insertion the other lever was followed by nothing. Trials were pseudorandomized such that no more than two of the same trial followed in sequence with intertrial intervals of approximately 2 min. Sign tracking sessions were roughly 1 hr in length. Thirty min prior to each sign-tracking session, intraperitoneal injections of clozapine-n-oxide (CNO) were given. CNO was dissolved in sterile water to a concentration of 0.001 g/ml, and was given at a relatively low 1 mg/kg dose that we and others have found effective for behavioral studies including sign-tracking tasks (Smith et al. 2016).

After the 12 sign-tracking sessions, tests of free feeding were given. Rats were given CNO injections as above, and then 30 min later given access to 16 g of grain pellets. Free feeding was conducted in 2 consecutive sessions 24 hrs apart within 5 days of the last sign-tracking session. Rats were given 1 hr to consume 16 g of grain pellets. Food weight was measured pre- and post-feeding to calculate grams consumed.

### 2.5 Histology

After completion of behavioral procedures, rats were deeply anesthetized with 1ml phenobarbital and perfused with 0.9% saline solution for approximately 6-8 min followed by perfusion of 10% formalin until fixture of head and neck tissue (approximately 3-4 min). Brain tissue was extracted, saturated with 20% sucrose and frozen to −80°C until sliced to 60 µm thick sections and mounted. Slides were coverslipped with Vectashield mounting medium containing DAPI (Vector Labs). Fluorescent expression was imaged via Olympus U-HGLGPS. Some animals (N = 5) underwent mCherry immunofluorescence if they showed dimmer viral expression than expected. For this, 60 µm slices were washed in 0.1M PBS (3×10 min), blocked in a 3% normal donkey serum (one hour), and incubated overnight in primary antibody (rabbit anti-DsRed, 1:500; Clontech). The next day, slices were again rinsed in 0.1M PBS (3×10 minutes) and then incubated for 4-5 hr in secondary antibody (donkey anti-rabbit Alexafluor 594, 1:500; Thermo Scientific). After a last rinse in 0.1M PB (3×10 min), slices were mounted and coverslipped with DAPA-containing Vectashield (Vector Labs). Per-animal expression was manually transcribed onto printed images (Paxinos & Watson, 2007) and then transcribed digitally via PowerPoint (Microsoft) at 90% transparency. Per-animal expression maps were then combined into group expression maps by digitally overlaying the expression areas. VP expression was defined by Paxinos and Watson (2009) coordinates (AP: −0.12, ML: 2.4, DV: −8.2) with the target being ventral to anterior commissure and lateral/anterior to the substantia innominata and lateral preoptic area but dorsal to magnocellular preoptic nucleus. NAcLSh expression was targeted on coordinates (AP: +1.44mm, ML: 2.0mm, DV: −8.2mm) with expression localized to the area lateral and ventral to the accumbens core but dorsal to the most rostral portion of VP and medial to the endopiriform nucleus.

### 2.6 Fluorescent Retrograde Tracer (CAV2-zsGreen)

As the GFP tag in the subset of rats with CAV2-Cre-GFP injections was not reliably detectable, we estimated CAV2 spread in 6 separate animals (3 received a 1.0µl injection into the VP; 3 received a 1.0µl injection into the NAcLSh) that were unilaterally injected with CAV2-zsGreen (Zweifel Laboratory, University of Washington). Injection, histology, and imaging procedures were performed as above.

### 2.7 Statistical Modeling and Analysis

All statistical tests were carried out using R (R Core Team, 2013). Categorical variables with multiple levels (e.g., Cue Block) were dummy coded to make predetermined comparisons between levels (e.g., pre CS+ vs. CS+ block, post CS+ vs. CS+ block). All linear mixed models are fit by maximum likelihood and t-tests use Satterthwaite approximations of degrees of freedom (R; “lmerTest”, Kuznetsova et al. 2015). The reported statistics will include parameter estimates (β values), confidence intervals (95% bootstrapped confidence intervals around dependent variable), standard error of the parameter estimate (SE), and p-values (R; “lmerTest”). Graphs were created through GraphPad Prism (version 7.0a) and designed with Adobe Illustrator.

Sign tracking data was analyzed for lever presses per minute (total lever presses over the session / minutes of lever availability) in a linear mixed model which accounts for the fixed effects of group (NAcLSh→VP inhibition group and NAcLSh→VP controls were analyzed in an individual model while the opposite projection group and its control was analyzed separately) by interaction with session of training (sessions 1-12), and for random effects of individual rat intercepts (i.e., individual session one values) and slopes (i.e., individual rat learning rates). See example below:

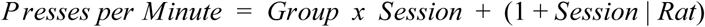

When trends appeared non-linear, transformations of session were tested to best fit learning rate curves (including logarithmic, quadratic, and exponential growth) and utilized if the transformation was of statistically better fit as determined by an analysis of variance comparing model deviances (p < 0.05, R; “anova” {lmerTest}) for nested models and Akaike information criterion (AIC) for non-nested model comparison (i.e., the model with the lowest AIC, ΔAIC = Model AIC - Null Model AIC). It is worth noting that when examining logarithmic or exponential components of session, the linear term is not included in the model. Both the logarithmic and exponential curves fit an increasing (or decreasing) function that levels off by themselves. They do not need (and should not add) the linear term as it is redundant. However, quadratic models include the linear fit as well, as in this case it is not redundant. An equation with 𝒳 and 𝒳^2^ in it is a second order polynomial. It fits empirical curves that are partly linear but have a bend or leveling off. The coefficient for the 𝒳 term shows how much the curve increases and the 𝒳2 term shows how prominent the bend is. Parameter estimates for the transformed independent variable (i.e. session) are presented in the units of the dependent variable (i.e. ppm) and can be interpreted such that the dependent variable units increase for every transformed component of the independent variable. For example, the logarithmic transformation of session might yield a significant effect with an estimate of 1.28 ppm and this can be interpreted to mean that the press rate increases 1.28 for every increase of one log session.

Free feeding data was similarly analyzed for amount of food consumed (g) in a linear mixed model which accounts for the group (i.e., NAcLSh→VP inhibition group and NAcLSh→VP control were analyzed in an individual model while the opposite projection group and its control were analyzed separately), individual animal weights taken during free feeding experiments, session (i.e., session 1 and 2 of free feeding), and random effects of rat.

## 3. Results

### 3.1 Viral expression

All animals were evaluated for robust expression of hM4Di receptors in the upstream area of interest (i.e. in VP for the VP→NAcLSh projection); those without clear expression or expression outside the area of interest were excluded from analysis (n = 15, due mainly to a faulty virus batch). Additionally, 3 animals were excluded during behavioral testing for health reasons. Thus, analysis was run with group sizes as: 14 animals in NAcLSh→VP analysis (control group = 6; inhibition group = 8), 19 animals in VP→NAcLSh analysis (control group = 9, inhibition group = 10). Linear mixed model analysis of repeated measures data can accommodate uneven group sizes (Gibbons et al. 2010).

The vast majority of animals in the NAcLSh→VP inhibition group exhibited hM4Di-mCherry expression in the lateral shell area as previously defined between bregma +1.20-2.16 mm with one animal showing some expression slightly more medial (although this animal had anterior expression more lateral; Fig.1A). Two animals had very discrete expression of hM4Di-mCherry in anterior portions of VP, but with the vast majority of expression in NAcLSh, and were thus included in analyses. Similarly, animals in the NAcLSh→VP site-specific control exhibited mCherry expression in the NAcLSh area. Notedly, mCherry expression from the control virus appeared more robust, and tended spread into a larger area than the

DREADD-containing inhibitory virus (Fig.1B). These controls were included in analysis given that they would control for viral mediated gene delivery at a level covering the DREADD expression areas and even beyond. Animals in the VP→NAcLSh inhibition group had robust hM4Di expression in the defined VP region between bregma −0.36-+0.36 mm (Fig.2A). Two animals showed more rostral VP expression up to +1.68 mm, but by far the most dense expression occurred in the VP in these animals. Given the minor spread beyond the target area of hM4Di expression, we refer to manipulations here as pathway-biased rather than pathway-specific. VP → NAcLSh control group expression was similarly greatest in VP with more spread dorsally as seen in the NAcLSh → VP control group (Fig. 2B).

**Figure 2:**
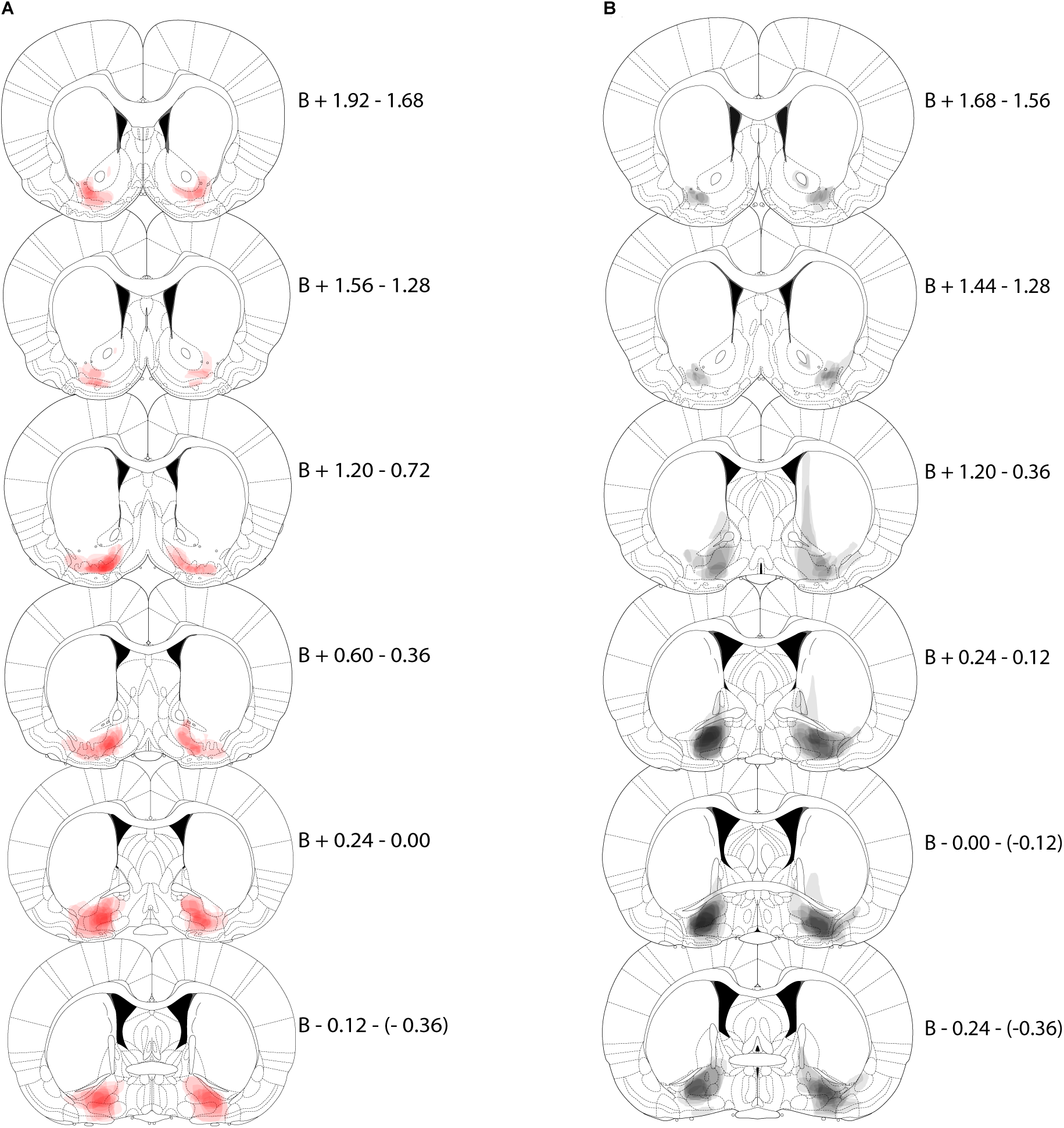
VP → NAcLSh inhibition histology. Each coronal section represents a range of three sections relative to Bregma (B, in mm). Expression for each animal is plotted at 90% transparency and then per-animal expression is overlaid. Sections run anterior (top) to posterior (bottom). A) Expression map of AAV-hSyn-DIO-hM4D(Gi)-mCherry in VP following CAV2-Cre injections in NAcLSh (i.e., VP → NAcLSh inhibition animals; red). B) Expression map of AAV-hSyn-DIO-mCherry in VP (i.e., controls; black).

Select animals in manipulation groups (NAcLSh→VP inhibition [n= 5 of 8], VP→NAcLSh inhibition [n = 3 of 10]), showed hM4Di-mCherry expression that could be characterized as “fleck-like” where expression was not clearly arranged in the neuronal membranes. This expression was only visible in the TexasRed filter, and not in other filters, nor was there any sign of cell death as judged by DAPI expression. This provided confidence that the expression was the mCherry tag, as did the statistically similar results on the main behavior of interest (CS+ responding) of these animals (Supplementary Fig. 1).

**Figure 1:**
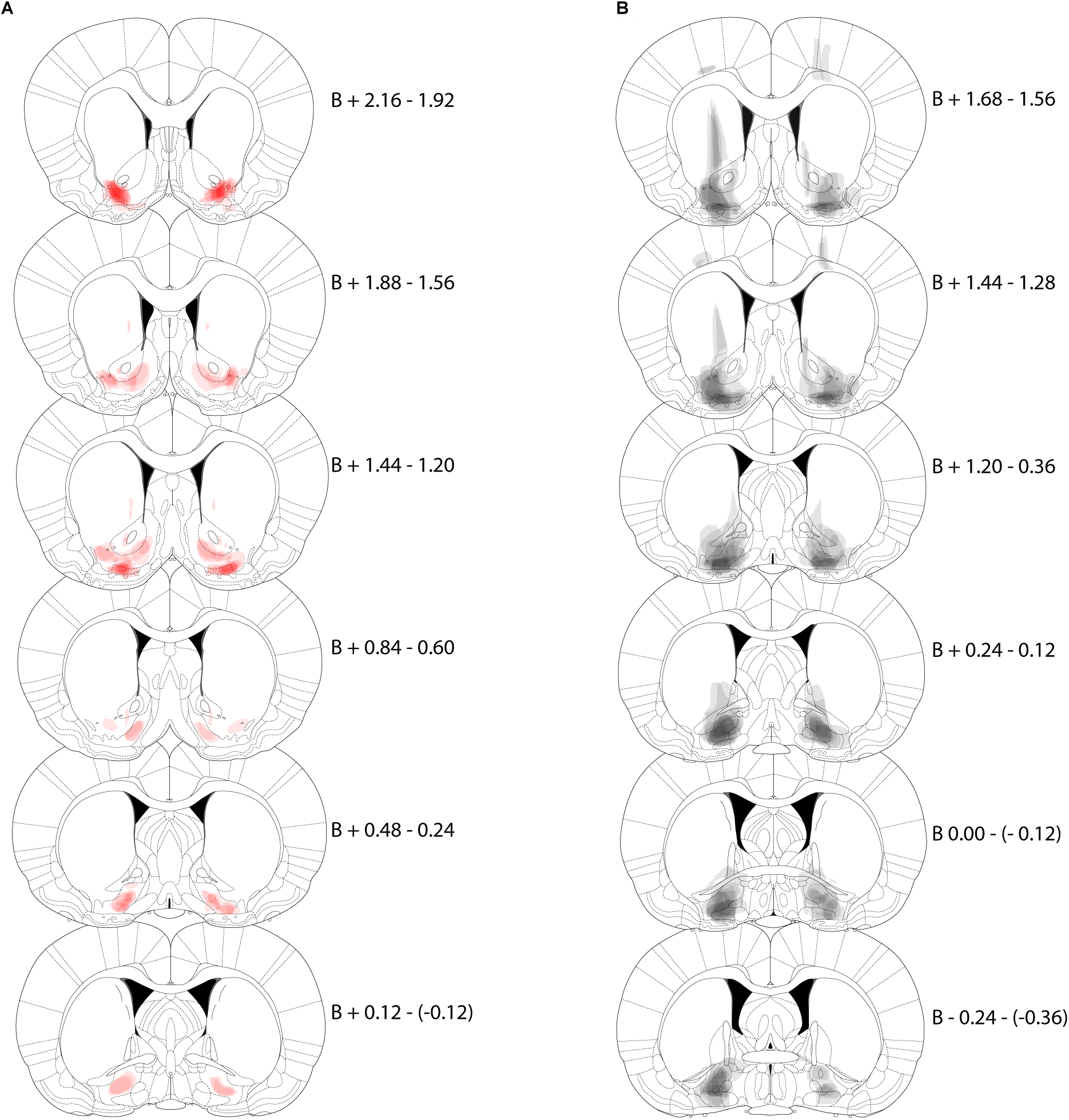
NAcLSh→VP expression maps. Each coronal section represents a range of three sections relative to Bregma (B, in mm). Expression for each animal is plotted at 90% transparency and then per-animal expression is overlaid. Sections run anterior (top) to posterior (bottom). A) Expression map of AAV-hSyn-DIO-hM4D(Gi)-mCherry in NAcLSh following CAV2-Cre injections in the VP (i.e., NAcLSh→VP inhibition animals; red). B) Expression map of AAV-hSyn-DIO-mCherry in NAcLSh (i.e., controls; black).

*CAV expression estimation (CAV2-zsGreen).* In the animals who received NAcLSh injections of CAV-zsGreen, discrete localization of zsGreen was seen in lateral VP (Fig. 3A). In animals who received VP injections, discrete localizations of zsGreen was seen in very lateral portions of our NAcLSh target (Fig. 3B). In both of these cases, the zsGreen expression was highly confined anatomically.

**Figure 3:**
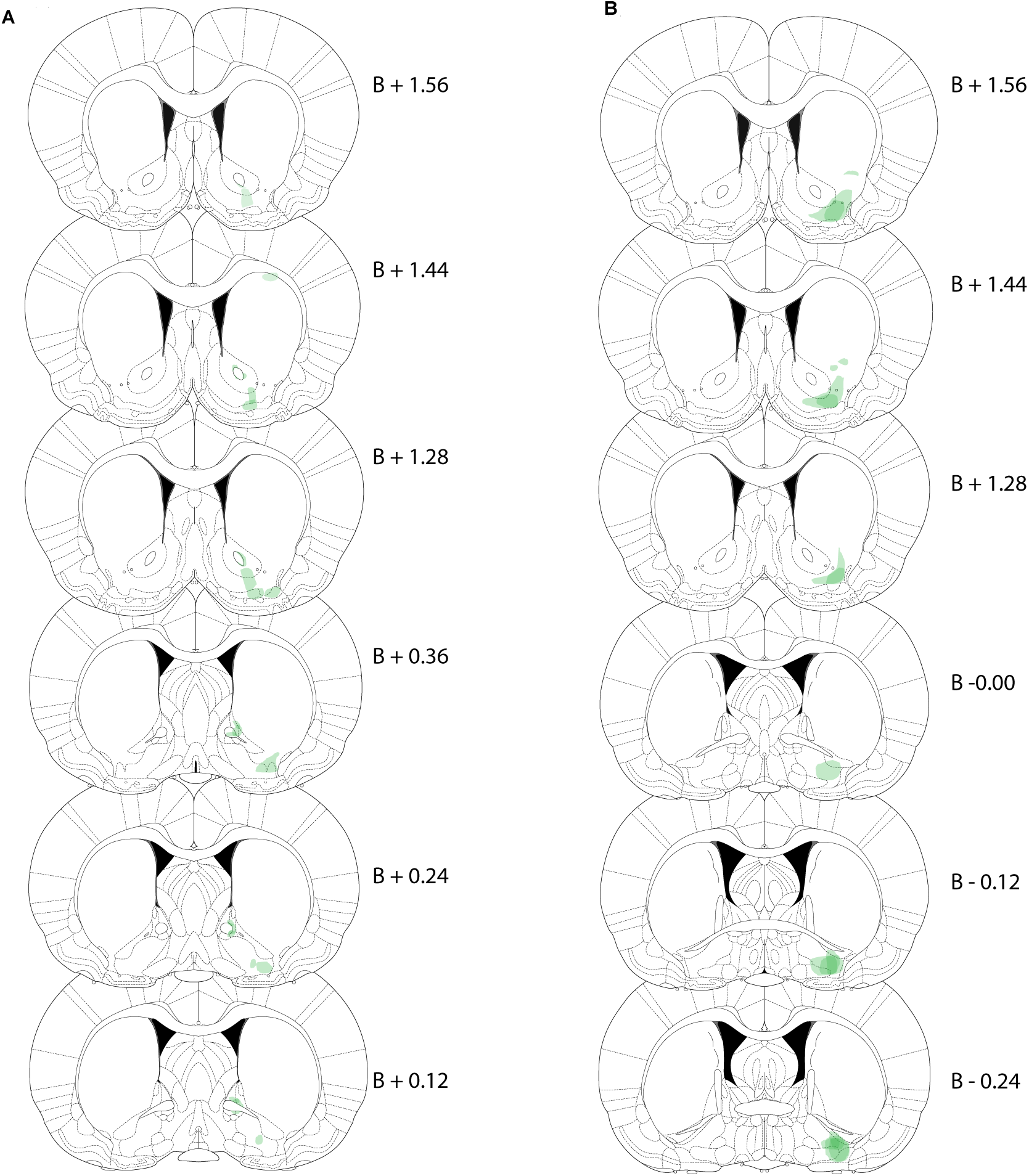
CAV-zsGreen histology. A) Expression of retrograde tracer CAV2-zsGreen after injection into NAcLSh target (transparency: 70%; green). B) Expression of CAV2-zsGreen after injection into the VP target (transparency: 70%; green). Expression will reflect retrograde transport into neurons projecting to the injection site, including interneurons.

### 3.2 NAcLSh to VP projection inhibition analysis

#### 3.2.1 CS+ presses per minute (ppm) during sign-tracking

The inhibition of the NAcLSh→VP pathway resulted in a decrease in ppm toward the CS+ lever compared to its control group. The NAcLSh→VP projection data of CS+ ppm over time appeared non-linear in form; comparison of a model containing quadratic and linear components against a model containing only linear components revealed a significant contribution of quadratic session to the model (𝒳^2^(4) = 72.62, p < 0.001). Thus, a linear mixed model was constructed with CS+ ppm by main effects of group (inhibition vs. control) by session (1 - 12; both linear and quadratic fits), with random effects of animal starting level and learning rate:

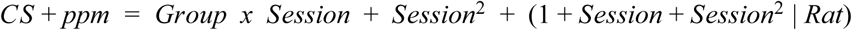

This model had both significant quadratic (est:-0.39 ppm; CI: −0.68-(−0.11); SE: 0.15; p = 0.017) and linear (est: 6.40 ppm; CI: 3.15-9.72; SE: 1.69; p = 0.002) fit components and did not show a main effect of group (est: −4.48 ppm; CI: −17.9-9.18; SE: 6.14; p = 0.478). The group by linear fit (est: 2.36 ppm; CI: 0.34-4.27; SE: 0.86; p = 0.015) interaction was significant. This result indicated that animals with the NAcLSh→VP pathway inhibited showed markedly reduced sign-tracking behavior as a function of training time compared to controls (Fig. 4A).

**Figure 4:**
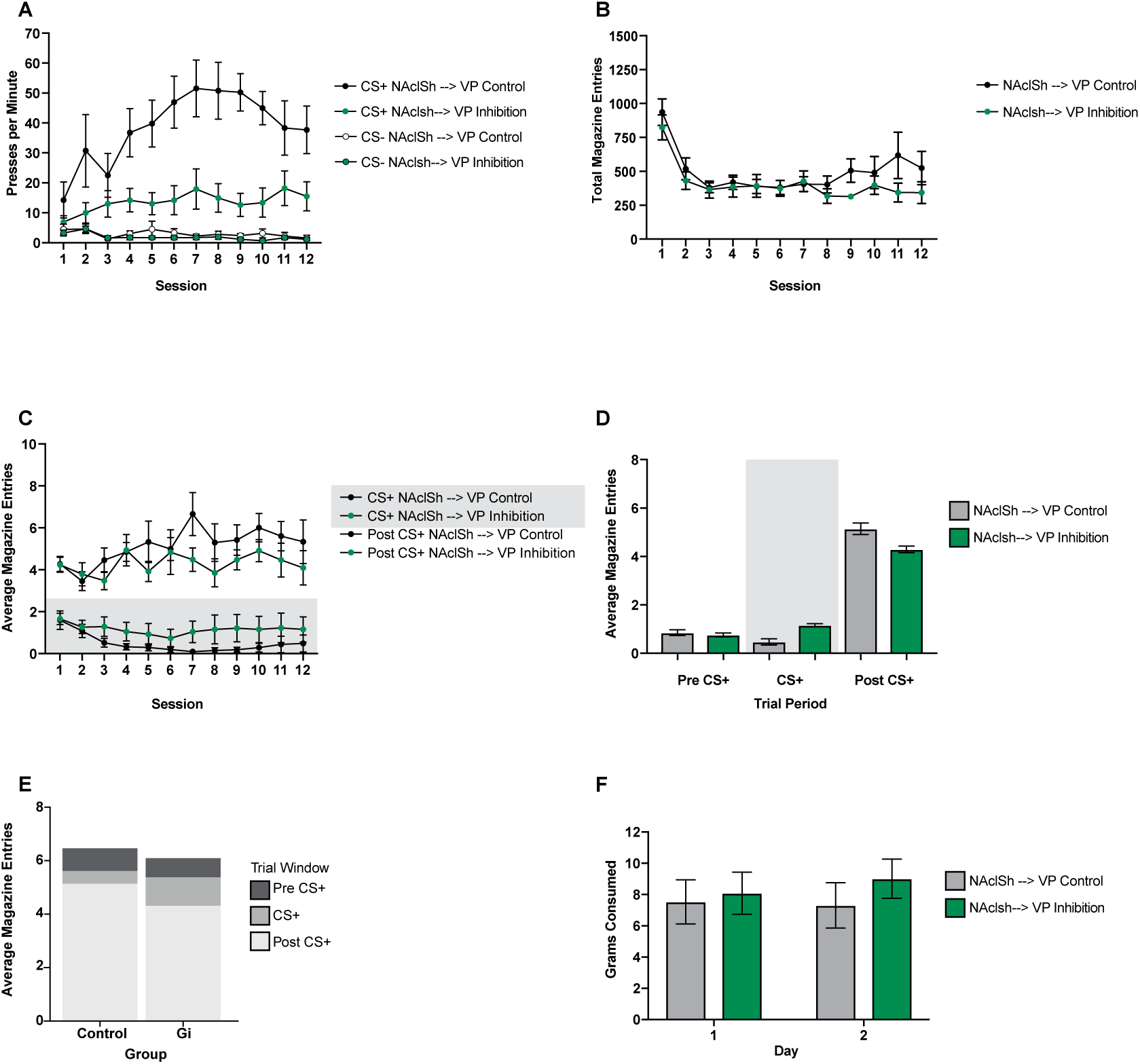
Effects of NAcLSh→VP inhibition. A) Presses per minute (ppm) on the CS+ lever over the 12 training sessions for the NAcLSh→VP inhibition group (green) and the NAcLSh→VP control group (black). B) Ppm on the CS- lever for both groups. C) Average magazine entries per session during the 10 sec CS+ presentation (shaded grey background) and the 10 sec post CS+ block (i.e., reward delivery) for both groups. D) Average magazine entries per 10 sec block type (10 sec Pre CS+, 10 sec CS+, 10 sec Post CS+) in the NAcLSh→VP inhibition group (green) and the NAcLSh → VP control group (grey). E) Average magazine entries per block type in the NAcLSh→VP control group (left) NAcLSh → VP inhibition group (right) with the whole bar representing the total magazine entries made during the 30 second trial period encompassing all three blocks. F) Grams of food consumed over the two free feeding sessions in NAcLSh→VP inhibition group (green) and the NAcLSh→VP control group (grey). For all graphs, bars and lines show mean and errors show +/- SEM.

#### 3.2.2 CS- ppm during sign-tracking

Both the NAcLSh→VP inhibition group and their control group decreased ppm toward the non-reinforced lever similarly. CS- ppm data appeared linear and a linear mixed model fitting CS- press rates by main effects of group by session (1-12; natural log transformation), with random effects of animal starting level and learning rate:

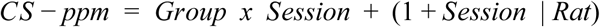

We detected no significant difference between Gi and controls (est: 0.80 ppm; CI: −1.35-3.26; SE: 1.12; p = 0.486) nor a significant interaction of group by session (est: 0.04 ppm; CI: −0.14-0.21; SE: 0.09; p = 0.676), indicating that groups did not differ overall in interaction with the non-predictive lever nor in how they terminated pressing overtime. There was a significant main effect of session (est: −0.19 ppm; CI: −0.28-(−0.09); SE: 0.04; p < 0.001), showing that all groups decreased CS- pressing over the course of training (Fig. 4A).

#### 3.2.3 CS+ vs. CS- ppm during sign-tracking summary

Ppm on the reinforced (CS+) and non-reinforced (CS-) lever were compared within each group by fixed effects of time (i.e., session) and lever type (CS+ vs. CS-). Both NAcLSh→VP inhibition animals and their control group showed a preference for the reinforced CS+ lever over the non-reinforced CS- lever. Although the degree of CS+ sign-tracking was reduced in the inhibition group compared to controls (see analysis above), the inhibition group still showed a preference for the CS+ compared to the CS-. In short, sign-tracking during NAcLSh→VP inhibition was reduced but was not eliminated to the level of CS- sign-tracking.

#### 3.2.4 NAcLSh→VP inhibition group

Data appeared linear and a linear mixed model was used to analyze press rate by lever type and session interaction with random slope and rat intercepts:

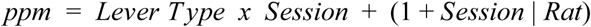

There was a significant main effect of press rates comparing the CS- to CS+ levers (est: 6.69 ppm; CI: 2.74-10.8; SE: 2.08; p = 0.002) with on average nearly 7 ppm greater towards the reinforced lever. There was not a significant main effect of session (est: 0.20 ppm; CI: −0.18-0.58; SE: 0.20; p = 0.346). However, a significant session by cue interaction (est: 0.81 ppm; CI: 0.24-1.36; SE: 0.28; p = 0.005) illustrated an increase in sign-tracking to the reinforced CS+ lever over time (Fig. 4A).

#### 3.2.5 NAcLSh→VP site-specific control group

A quadratic transformation in session significantly contributed to the model (𝒳^2^(4) = 22.40, p < 0.001) and thus was included in the final model:

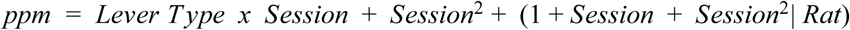

There was a significant main effect of press rates toward the predictive lever (est: 22.1 ppm; CI: 14.0-30.6; SE: 4.20; p < 0.001). Both linear (est: 5.84 ppm; CI: 2.61-8.96; SE: 1.62; p = 0.006) and quadratic (est: −0.39 ppm; CI: −0.67-(−0.11); SE: 0.14; p = 0.029) transformations in session were significant. A linear session by cue interaction (est: 1.94 ppm; CI: 0.73-3.01; SE: 0.57; p < 0.001) showed a significant increase in presses toward the CS+ lever over sessions compared to the CS- lever (Fig. 4A).

#### 3.2.6 Magazine entries

Groups did not differ in their overall magazine entries per day nor in how they entered the magazine over sessions. Total magazine entries per day appeared linear and were analyzed by a linear mixed model with fixed effects of group by session and quadratic fit of session with random slopes and intercepts:

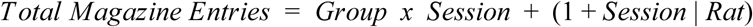

The main effect of group was not significant (est: −31.4 entries; CI: −226-176; SE: 104; p = 0.767). The main effect of session was significant (est: −16.1 entries; CI: −30.5-(−1.95); SE: 7.51; p = 0.049), indicating that all animals decreased their number of entries by the end of training. The interaction of group by session was not significant (est: 15.8 entries; CI: −14.6-46.7; SE: 15.0; p = 0.311), and thus groups did not differ over sessions in their magazine entry behavior (Fig. 4B).

#### 3.2.7 Magazine entries by cue block

Although the NAcLSh→VP inhibition group and their control group did not differ in overall magazine entries, as above, there was a difference in how each group distributed their entries with respect to the CS+ versus post-CS+ time blocks over sessions. The control group displayed more traditional sign-tracking behavior where they developed a decreased magazine entry rate during cue presentation and increased magazine entries during the post-cue block when reward was available. The inhibition group did not show this shift over time, exhibiting less of a difference in entries between the CS+ and post CS+ block.

Magazine entries recorded during ten-sec cue blocks (pre CS+, CS+, post CS+ [i.e. reward delivery]) each were averaged over 25 trials per day (i.e., an average entries per trial in the pre CS+ block). A predetermined contrast analyzed the CS+ presentation block against the reward delivery block and if more magazine entries were performed during the reward block (as expected) compared to the cue presentation. Another contrast analyzed the 10 second block prior to CS+ presentation against the reward delivery and if magazine entries were greater during the reward (as expected) compared to the time prior to cue delivery. The average entries per block appeared linear and were analyzed in a linear mixed model by fixed effects of session, group, and cue, with random effects for individual rat intercepts (random slopes could not be included due to failed convergence):

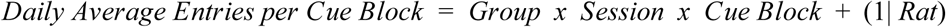

No significant main effect of group was found (est: −0.02 avg. entries; CI: −0.90-1.03; SE: 0.49; p = 0.961). However, a significant session effect was found (est: 0.10 avg. entries; CI: 0.05-0.15; SE: 0.03; p < 0.001). Regardless of group, contrasts to compare pre CS+ to post CS+ blocks was significant with 2.85 average entries greater during the post CS+ block compared to pre CS+ (est: −2.85 avg. entries; CI: −3.42-(−2.32); SE: 0.29; p < 0.001). CS+ presentation to post CS+ block was also significant with 2.90 average entries greater during the post CS+ block compared to pre CS+ (est: −2.09 avg. entries; CI: −3.56-(−2.36); SE: 0.29; p < 0.001). The trend towards greater entries during the post CS+ block developed over time as seen in a significant pre CS+ to post CS+ by session interaction (est: −0.16 avg. entries; CI: −0.22-(−0.08); SE: 0.04; p < 0.001) and significant CS+ to post CS+ by session interaction (est: −0.15 avg. entries; CI: −0.21-(−0.07); SE: 0.04; p < 0.001; Fig. 4C). Groups differed in how they entered the magazine over sessions, with a significant group by session interaction (est: 0.13 avg. entries; CI: 0.02-0.22; SE: 0.05; p = 0.018). A three way interaction of group by session by CS+ vs. post CS+ blocks was significant (est: −0.20 avg. entries; CI: −0.33-(−0.04); SE: 0.08; p = 0.009), indicating that the control group made slightly more magazine entries during the post-CS+ reward block, and fewer entries during the CS+ itself, over sessions compared to the inhibition group (Fig. 4C). Notedly, this difference was only seen over sessions and was not present overall as determined by a non-significant CS+ versus post CS+ by group interaction (est: −0.17 avg. entries; CI: −1.29-0.91; SE: 0.58; p = 0.767; Fig. 4D, 4E). Groups did not differ over sessions in how they entered the magazine during the ten second blocks before vs. during CS+ presentation (i.e., the three way interaction of group by session by contrast of pre CS+ to post CS+; est: −0.10 avg. entries; CI: −0.25-0.05; SE: 0.08; p = 0.173). The overall interaction of preCS+ versus post CS+ by group interaction was not significant (est: −0.15 avg. entries; CI: −1.16-1.03; SE: 0.58; p = 0.942; Fig. 4D, 4E).

#### 3.2.8 Free feeding

The NAcLSh→VP inhibition and the control groups did not differ in the amount of food consumed during free feeding tests nor did their weights impact feeding behavior. A linear mixed model of total grams of food consumed as a function of group (control versus inhibition) by session (sessions 1 and 2) and body weight (in grams) by session, with random effects of individual animal, was constructed:

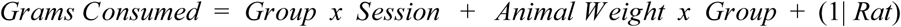

There was not a significant main effect grams consumed by session of testing (est: 0.42 grams; CI: −0.44-1.18; SE: 0.41; p = 0.319) nor main effect of grams consumed by group (est: −11.25 grams; CI: −32.8-9.79; SE: 11.1; p = 0.325). Grams consumed did not differ by body weight (est: −0.01 grams; CI: −0.05-0.02; SE: 0.02; p = 0.410). There was not a significant group by session interaction (est: −1.33 grams; CI: −3.23-0.26; SE: 0.82; p = 0.123) nor group by weight interaction (est: 0.03 grams; CI: −0.03-0.10; SE: 0.03; p = 0.323; Fig. 4F).

### 3.3 VP→NAcLSh projection inhibition analysis

#### 3.3.1 CS+ ppm during sign-tracking

The VP→NAcLSh inhibition group and their controls did not differ in sign-tracking to the CS+ lever. The VP→NAcLSh projection data of CS+ responses over time appeared non-linear in form and model fit significantly improved after addition of a quadratic fit of session (𝒳^2^(4) = 50.9, p < 0.001) and thus were included in the final model. The same model structure for CS+ ppm analysis for the opposite projection group (see above) was used here:

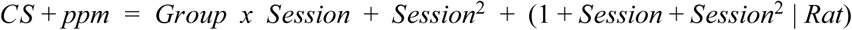

Results showed an insignificant main effect of group (est: −4.57 ppm; CI: −12.5-4.02; SE: 4.16; p = 0.285). A significant linear component of session (est: 3.16 ppm; CI: 0.95-5.45; SE: 1.13; p = 0.012) identified that all animals learned over training. The quadratic component of session was also significant (est: −0.176 ppm; CI: −0.34-(−0.01); p = 0.037) again indicating that animals learned to acquire CS+ responding at a similar rate. Group by linear session was not significant in analysis (est: 0.213 ppm; CI: −0.93-1.40; SE: 0.56; p = 0.705; Fig. 5A), indicating that all groups learned at the same rate over sessions. Thus, unlike the reduced sign-tracking observed with NAcLSh→VP inhibition compared to their controls, sign-tracking with VP→NAcLSh inhibition appeared equivalent to their controls. Sign-tracking levels were generally lower in the VP→NAcLSh groups compared to those targeting the NAcLSh→VP, but still in a normal range based on prior studies, which highlights the importance of controls that are specific to pathways being targeted (Fig. 5A).

**Figure 5:**
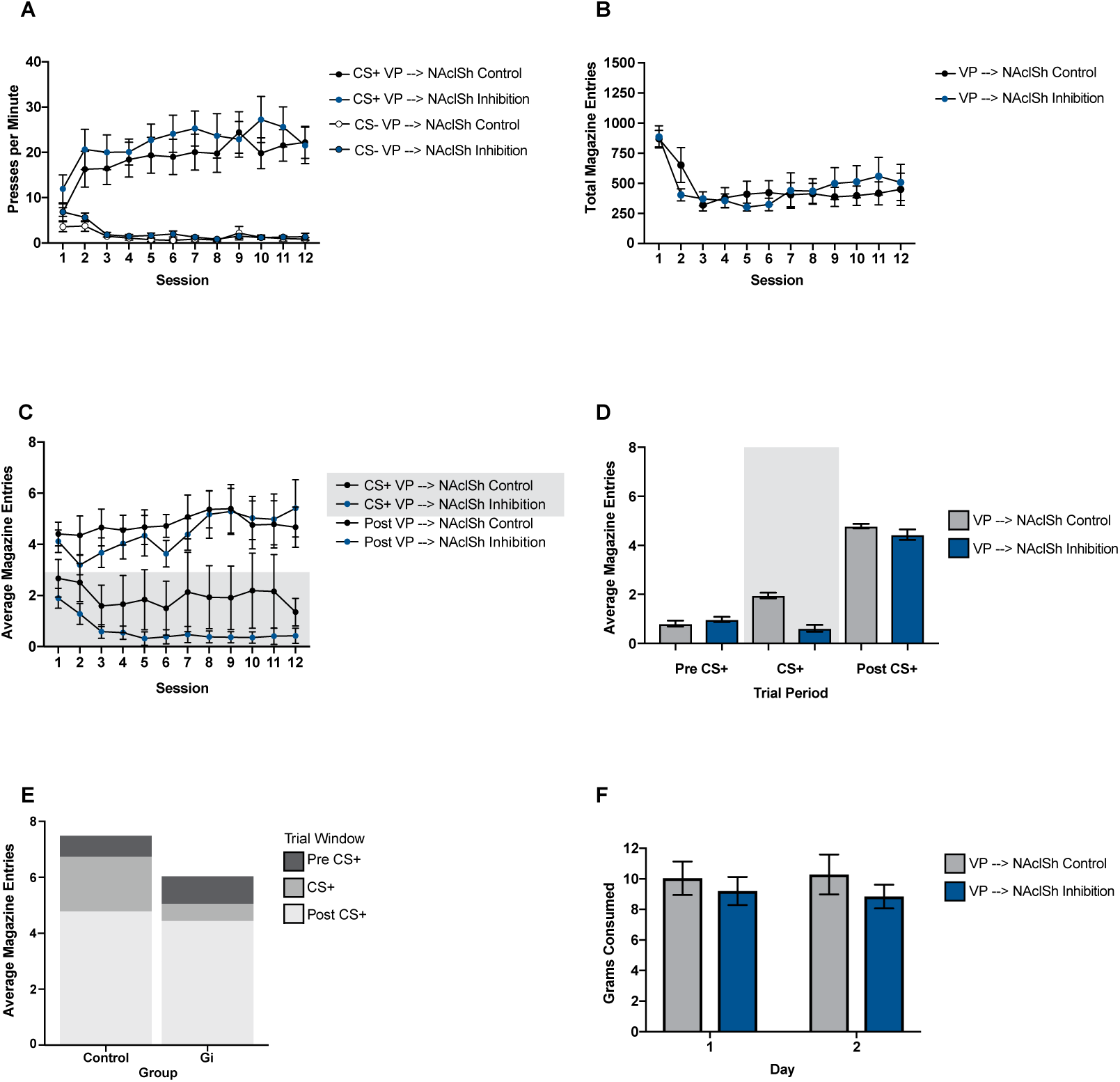
Effects of VP → NAcLSh inhibition. A) Presses per minute (ppm) on the CS+ lever over the 12 training sessions for the VP → NAcLSh inhibition group (blue) and the VP → NAcLSh control group (black). B) Ppm on the CS- lever for both groups. C) Average magazine entries per session during the 10 sec CS+ presentation (shaded grey background) and the 10 sec post CS+ block (i.e., reward delivery) for both groups. D) Average magazine entries per 10 sec block type (10 sec Pre CS+, 10 sec CS+, 10 sec Post CS+) in the VP → NAcLSh inhibition group (blue) and the VP → NAcLSh control group (grey). E) Average magazine entries per block type in the VP → NAcLSh control group (left) VP → NAcLSh inhibition group (right) with the whole bar representing the total magazine entries made during the 30 second trial period encompassing all three blocks. F) Grams of food consumed over the two free feeding sessions in VP → NAcLSh inhibition group (blue) and the VP → NAcLSh control group (grey). For all graphs, bars and lines show mean and errors show +/- SEM.

#### 3.3.2 CS- ppm during sign-tracking

A significant main effect of group showed that rats with VP→NAcLSh inhibition pressed the non-reinforced CS- more than their controls. However, over sessions both groups decreased press rates similarly toward the CS-. Data appeared linear and the model of CS- responses by fixed effects of group by linear session was constructed:

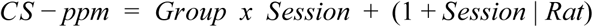

There was a significant main effect of group (est: −2.02 ppm; CI: −3.79-(−0.34); SE: 0.88; p = 0.034). There was a significant main effect of linear session (est: −0.27 ppm; CI: −0.38-(−0.17); SE: 0.05; p < 0.001), indicating that all animals decreased pressing toward the CS- lever. The group by linear session interaction was not significant (est: 0.19 ppm; CI: 0.01-0.40; SE: 0.10; p = 0.063; Fig. 5A), indicating that all animals showed similar decreases in CS- press rates over sessions.

#### 3.3.3 CS+ vs. CS- ppm during sign-tracking

Both the VP→NAcLSh inhibition group and their control group prefered the reinforced CS+ lever over the non-reinforced CS- lever. Press rates on the CS+ and CS- levers were compared within each group by fixed effects of time (i.e. session) and lever type (CS+ v. CS-). The following model structure is used in the next two sections.

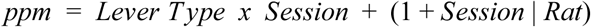

#### 3.3.4 VP→NAcLSh inhibition group

The VP→NAcLSh inhibition group prefered the CS+ lever over the CS- lever. There was a significant main effect of cue type (est: 12.4 ppm; CI: 8.24-16.6; SE: 2.11; p < 0.001) such that the CS+ was significantly preferred over the CS- in pressing behavior. There was not a significant main effect of linear session (est: 0.20 ppm; CI: −0.32-0.74; SE: 0.26; p = 0.450).

However, a lever cue by session interaction was significant (est: 1.14 ppm; CI: 0.58-1.69; SE: 0.29; p < 0.001), such that rates of preference toward the CS+ lever increased over time (Fig. 5A).

#### 3.3.5 VP→NAcLSh site specific control group

The site specific control group also prefered the CS+ lever over the CS- lever. There was a main effect of cue type (est: 9.73 ppm; CI: 6.10-13.4; SE: 1.78; p < 0.001) as confirmation of this. There was a significant main effect of session (est: 0.39 ppm; CI: 0.14-0.67; SE: 0.14; p

= 0.018) and session by leve type interaction (est: 1.13 ppm; CI: 0.65-1.64; SE: 0.24; p < 0.001), indicating that control animals interacting with the CS+ lever more over time (Fig. 5A).

#### 3.3.6 Magazine entries

Total magazine entries per session appeared linear and model fit a linear mixed model of magazine entries by group and day with random slopes and intercepts was constructed:

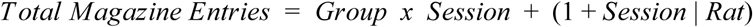

The VP→NAcLSh inhibition and site specific control groups did not differ in magazine entries as determined by a non-significant group effect (est: 122 entries; CI: −50.8-296; SE: 91.4; p = 0.198). Magazine entries did not change over sessions as seen in an insignificant main effect of session (est: −11.7 entries; CI: −30.0-7.27; SE: 8.76; p = 0.196). The groups similarly maintained magazine entry rates over time as seen in an insignificant group by session interaction (est: −19.7 entries; CI: −57.3-12.8; SE: 17.5; p = 0.276; Fig. 5B).

#### 3.3.7 Magazine entries during cue blocks

Despite the lack of effect of VP→NAcLSh inhibition on sign-tracking behavior and overall magazine entries, it did lead to a difference in magazine entry distribution: the VP→NAcLSh inhibition group made fewer entries in the CS+ presentation block and more entries in the post CS+ block compared to controls over sessions. The statistical model structure is the same as the opposite projection group above and includes the same cue block contrasts described above.

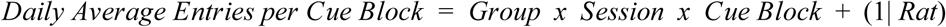

There was a significant main effect of group (est: 1.63 entries; CI: 0.41-2.95; SE: 0.66; p = 0.016) and linear session (est: 0.13 entries; CI: 0.06-0.20; SE: 0.03; p < 0.001). Main effects of pre-CS+ versus post-CS+ block (est: −2.64 entries; CI: −3.37-(−1.85); SE: 0.36; p < 0.001) and CS+ versus post-CS+ block (est: −2.27 entries; CI: −2.95-(−1.52); SE: 0.36; p < 0.001) were significant in that magazine entries were greatest in the post-CS+ reward block compared to the pre-CS+ and CS+ blocks (Fig. 5C, 5D). A group by linear session interaction was significant (est: −0.21 entries; CI: −0.33-(−0.07); SE: 0.07; p = 0.002). The group by pre-CS+ versus post-CS+ block interaction was not significant (est: −1.16 entries; CI: −2.48-0.23; SE: 0.72; p = 0.108; Fig. 5D). The group by CS+ versus post-CS+ block interaction was not significant (est: −0.94 entries; CI: −2.29-0.63; SE: 0.72; p = 0.193). Thus all animals tended to increase magazine entries over sessions during the reward block compared to the pre-CS+ block (est: −0.17 entries; CI: −0.27-(−0.07); SE: 0.05; p < 0.001) and compared to the CS+ presentation (est: −0.18; CI: −0.27-(−0.09); SE: 0.05; p < 0.001; Fig. 5C). The three way interaction of group by session by pre-CS+ vs. reward block was not significant (est: 0.10 entries; CI: −0.08-0.28; SE: 0.10; p = 0.307), indicating that all animals distributed their magazine entries similarly toward the post-CS+ reward block compared to the pre-CS+ block over sessions (Fig. 5C). The three-way interaction of group by session by CS+ presentation vs. reward block was significant (est: 0.33 entries; CI: 0.14-0.51; SE: 0.09; p < 0.001), indicating that over time, the VP→NAcLSh inhibition group entered the magazine less during the CS+ presentation and more during the post-CS+ reward block compared to controls.

#### 3.3.8 Free feeding

There was no difference in food consumption during free feeding tests between animals with VP→NAcLSh inhibition and their controls. The same model structure used in the NAcLSh→VP feeding analysis was used here:

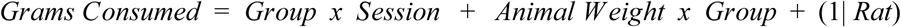

There was not a main effect of group (est: −7.48 grams; CI: −21.8-7.84; SE: 7.89; p = 0.354), meaning all animals consumed food similarly in free feeding tests. Animals remained consistent with their food intake over the two days of testing as seen in an insignificant main effect of session (est: 0.11 grams; CI: −1.10; SE: 0.61; p = 0.856) and insignificant main effect of group by session (est: −0.98 grams; CI: −3.61-1.53; SE: 1.23; p = 0.432). However, intriguingly, there was a main effect of weight (est: −0.028 grams; CI: −0.05-(−0.01); SE: 0.01; p = 0.020) whereby animals of greater weight tended to eat slightly less. However, this effect of weight was generalized over both groups as seen in an insignificant group by weight interaction (est: 0.022 grams; CI: −0.02-0.06; SE: 0.02; p = 0.322; Fig. 5F).

## 4 Discussion

To investigate the importance of NAcLsh and VP connections in the ability of reward cues to draw in motivated behavior, we used a pathway-biased chemogenetic manipulation strategy to inhibit projections from the NAcLSh to the VP, and vice versa, in a sign-tracking task. Inhibition of the NAcLsh→VP pathway resulted in a marked reduction of sign-tracking behavior. In stark contrast, inhibition of the reverse VP→NAcLSh pathway left sign-tracking behavior normal. However, it was not as though VP→NAcLSh inhibition was unremarkable. Those animals showed a shift over time to having greater magazine-directed behavior when reward was available after the post CS+ reward block versus to when the CS+ was presented compared to controls. Inhibition of the NAcLSh→VP instead had a greater tendency to exhibit magazine entries during the CS+ itself.

None of the sign-tracking or magazine entry effects we observed could be explained by a change in the motivational value of the food reward because free feeding behavior was unaffected by any manipulation. We hesitate to conclude that neither pathway is necessary for the motivation to eat, given that robust manipulations of either the NAcLSh or the VP can indeed affect eating behavior (e.g., Zhang and Kelley 2000; Cromwell and Berridge 1993). Two possibilities are either that the DREADD manipulation was subtle enough to leave other brain areas capable of driving eating behavior normally, or else that neither pathway is actually involved in free eating. In either case, the roles for other areas connected with the NAc and VP that are known to regulate eating (e.g., the lateral hypothalamus) deserve a similar circuit-based investigation. NAc projections to the lateral hypothalamus may well be an important circuit for feeding behavior, raising the intriguing possibility that similar manipulation of this pathway could reduce eating but not sign-tracking.

These results for magazine entries and sign-tracking further underscore the importance of NAc/VP interactions in regulating motivated behaviors, and give particular new importance to the NAcLSh→VP pathway in determining how motivationally attractive a reward-paired cue is to animals. It remains to be tested whether connections from other subregions of the NAc to the VP play a similarly important role. The NAc core and medial shell subregions participate importantly in sign-tracking behavior (Day and Carelli 2007; Flagel et al. 2011; Cardinal et al. 2002; Singer et al. 2016; Chang et al. 2012). One might thus suspect that any NAc output to the VP contributes similarly to incentive salience processes. As a potential caveat to this notion, we have recently found that a disconnection procedure to reduce communication between the NAc medial shell and VP not only failed to reduce sign-tracking, but instead increased it (Chang et al. 2018). This result was curious with respect to other studies that have shown decreases in cue-directed reward seeking and reward consumption using similar manipulations (e.g., Smith and Berridge 2007, Leung and Balleine 2015). Such findings collectively raise the importance of comparing NAc-VP subregional interactions across multiple motivation measures, and for comparing disconnection vs. pathway-based manipulation procedures, in order to fully elucidate how NAc and VP subregions interact for motivated behavior.

Another intriguing notion that these results raise is that the NAcLSh neurons that project to the VP do not appear to require a fully intact feedback signal in order to affect motivation. On the conceptual side, feedback projections are a common organizational principle of the brain and it remains unclear to what extent such feedback is necessary for the function of the principle site. In the case here, we can speculate that the motivationally-relevant information conveyed form the NAcLSh to the VP does not necessarily require information to be received back from the VP to the NAcLSh. Mechanistically, there are many details yet to resolve. For instance, it is possible that the VP projections to the NAcLSh do not actually synapse on the same NAcLSh neurons that project to the VP, which would help explain the pathway dissociation here. More generally, both the NAc and VP are composed of heterogeneous cell types (Smith et al. 2009; Root et al. 2015; Bobadilla et al 2017; Meredith and Totterdell 1999; Yang et al. 2018), suggesting that feedforward or feedback signals between these structures will likely result in a complex mix of inhibitory and excitatory neuronal effects (Hakan et al. 1992; Hakan et al. 1994; Hakan et al. 1995; Heimer & Wilson 1975; Churchill & Kalivas 1994; Napier & Mitrovic 1999). As such, the VP→NAcLSh DREADD manipulation here might not have led to the “relevant” sort of modulation of NAcLSh activity - whether targeting the relevant neuronal subpopulations or engaging the related activity patterns - to effect sign-tracking behavior (Gore et al. 2015). Nevertheless, the dissociation of NAc→VP inhibition reducing sign-tracking and shifting magazine-directed behavior to the CS+ time, and VP→NAc inhibition shifting magazine-directed behavior to the reward when it was presented, lays the groundwork for considering that the function of these forward and back projections might well be different for directing motivated behaviors.

## 5 Funding

This work was supported by funding from The Whitehall Foundation to KSS (2014-05-77) and the NIH (R01DA044199). The authors have no competing interests to declare.

## Acknowledgements

We would like to thank Alex Brown for assistance with histology.

## Supplemental Analysis & Figures

### “Fleck-like” expression analysis

Further analysis was used to determine if the animals within the NAcLSh→VP projection inhibition group that displayed atypical mCherry expression (n = 5) differed from the more cellular mCherry expressing animals in that group (n = 3). A linear mixed model of CS+ response rate, the main variable of interest, by histology outcome (fleck-like or cellular) by interaction of linear session and quadratic fit of session with random slopes and intercepts was constructed. This model structure is the same as used for the main analyses.

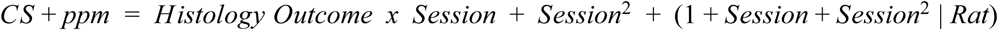

Data appeared non-linear and a quadratic fit of session significantly contributed to model fit (𝒳^2^(4) = 19.83, p < 0.001). Analysis determined no significant difference between the fleck-like animals and typically expressing animals in their CS+ ppm as seen in an insignificant main effect of histology outcome (est: 8.78 ppm; CI: −0.60-17.2; SE: 4.34; p = 0.077). Animals did not differ over sessions in both linear fit of session (est: 2.01 ppm; CI: −1.18-4.90; SE: 1.54; p = 0.226) and quadratic fit of session (est: −0.12 ppm; CI: −0.28-0.08; SE: 0.09; p = 0.273). Fleck-like and cellular expression animals did not differ over time in press rates as determined by an insignificant interaction of histology outcome and session (est: 0.06 ppm; CI: −1.15-1.18; SE: 0.53; p = 0.910).

The same analysis as above was performed on the fleck-like (n = 2) and cellular (n = 8) animals in the VP→NAcLSh projection inhibition group. Data appeared non-linear and model fit significantly improved upon addition of a quadratic fit of session (𝒳^2^(4) = 29.17, p < 0.001). An insignificant main effect of histology outcome (est: 0.07 ppm; CI: −13.2-15.5; SE: 7.08; p = 0.992) indicates that fleck-like animals did not differ from cellular expression animals in CS+ ppm. All animals maintained similar rates of pressing as determined by insignificant main effects of linear session (est: 3.07 ppm; CI: −1.07-7.30; SE: 2.03; p = 0.161) and quadratic fit of session (est: −0.19 ppm; CI: −0.50-0.08; SE: 0.14; p = 0.203). Fleck and cellular animals did not differ over sessions as seen by a non-significant histology outcome by session interaction (est: 0.53 ppm; CI: −1.93-2.91; SE: 1.16; p = 0.658).

---

**Supplemental Figure 1:**
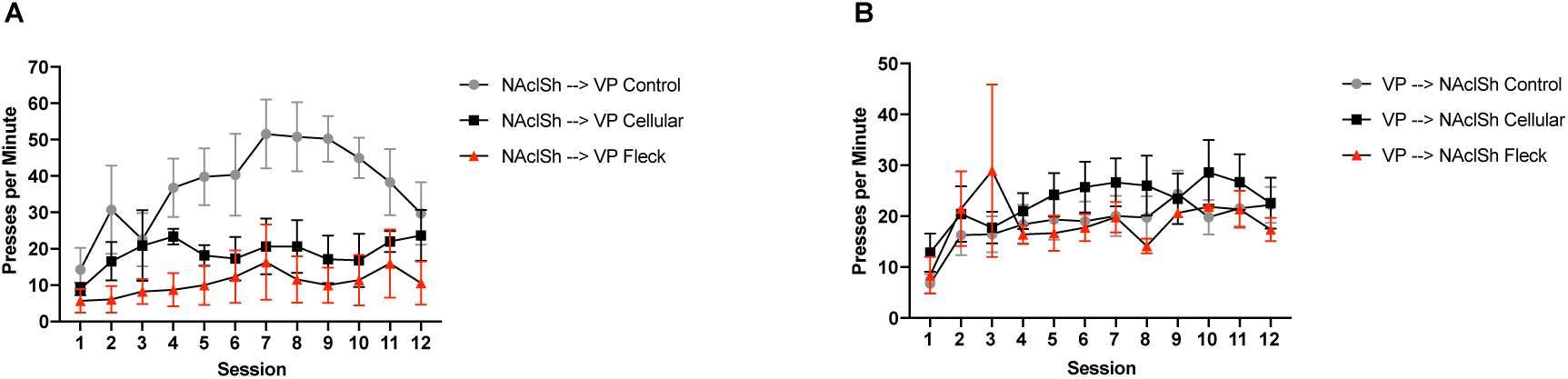
Fleck-like cellular expression versus typical cellular expression. A) Presses per minute (ppm) on the CS+ lever over the 12 training sessions in the sign-tracking paradigm in the NAcLSh→VP control group (n = 6, grey circle), inhibition group (n = 3, black square), and the NAcLSh→VP fleck-like expression group (n = 5, red triangle). B) Ppm on the CS+ lever over the 12 training sessions in the VP→NAcLSh inhibition group (n = 7, black square), the VP→NAcLSh control group (n = 9, grey circle), and the VP→NAcLSh fleck-like expression group (n = 3, red triangle). Lines denote mean and error bars denote +/- SEM.

